# Unbiased Atomistic Insight in the Mechanisms and Solvent Role for Globular Protein Dimer Dissociation

**DOI:** 10.1101/442889

**Authors:** Faidon Z. Brotzakis, Peter G. Bolhuis

## Abstract

Association and dissociation of proteins are fundamental processes in nature. While this process is simple to understand conceptually, the details of the underlying mechanism and role of the solvent are poorly understood. Here we investigate the mechanism and solvent role for the dissociation of the hydrophilic *β*-lactoglobulin dimer by employing transition path sampling. Analysis of the sampled path ensembles indicates that dissociation (and association) occurs via a variety of mechanisms: 1) a direct aligned dissociation 2) a hopping and rebinding transition followed by unbinding 3) a sliding transition before unbinding. Reaction coordinate and transition state analysis predicts that, besides native contact and vicinity salt-bridge interactions, solvent degrees of freedom play an important role in the dissociation process. Analysis of the structure and dynamics of the solvent molecules reveals that the dry native interface induces enhanced populations of both disordered hydration water and hydration water with higher tetrahedrality, mainly nearby hydrophobic residues. Bridging waters, hydrogen bonded to both proteins, support contacts, and exhibit a faster decay and reorientation dynamics in the transition state than in the native state interface, which renders the proteins more mobile and assists in rebinding. While not exhaustive, our sampling of rare unbiased reactive molecular dynamics trajectories shows in full detail how proteins can dissociate via complex pathways including (multiple) rebinding events. The atomistic insight obtained assists in further understanding and control of the dynamics of protein-protein interaction including the role of solvent.

PACS numbers:

## INTRODUCTION

Protein association and dissociation is essential for biologically relevant processes, such as cell signalling, DNA replication/transcription, cellular transport, immune response, gene editing [1], as well as for protein aggregation and self-assembly into structures with desired properties, e.g in food, colloids [2, 3]. Moreover, knowledge of the kinetics and mechanisms of association is crucial for understanding and controlling biochemical network and cascade reactions of processive or distributive nature [4, 5]. Yet, this kinetics is poorly understood even on the dimer level, and varies with the nature of the proteins[6]. While hydrophobic association/dissociation occurs through the dewetting effect [7, 8], association of hydrophilic proteins involves wet dimer native interfaces [9, 10], and has received significantly less attention, even though 70% of the protein-protein interfacial residues are hydrophilic [11]. The widely studied hydrophilic dimers Barnase-Barstar or Acetylcholinesterase-Fasciculin associate through a diffusion-limited reaction where the slow step is finding the transient encounter complex, a process accelerated by water assisted electrostatic steering between the charged hydrophilic interfaces [12–22]. However, little is known about hydrophilic dimers whose association is slower and not in the electrostatic steering regime, including many small proteins [23]. Several theoretical models stress the importance of water mediated interactions, assisting binding in the absence of steering. Ben Naim [24] proposed that hydrogen bond bridging water interactions between the two proteins are maximized towards the native dimer. Northrup et. al. [23] state that water assists binding through stabilization of a diffusion encounter complex which increases the rebinding probability. Association via rebinding increases for stronger isotropic (dispersion) interactions, which smooth the rugged energy landscape due to anisotropic (charged) interactions [25, 26]. In fact, water could also play such a smoothing role, e. g. by screening anisotropic salt bridges at the interface.

Various experimental studies on globular dimeric proteins, including NMR, PRE and sedimentation experiments [10, 14, 27–29], yielded (indirect) information on stable states, potential transition states and intermediates. However, atomistic details on association/dissociation pathways allowing direct insight into the mechanisms of dimer formation are still lacking. While molecular dynamics (MD) simulation can in principle provide such detailed insight, straightforward MD using all atom force fields is impractical as timescales of dissociation and association are on the order of milliseconds to seconds. Only recently a handful of MD studies addressed hydrophilic dimers (mostly on Barnase-Barstar) [12, 17, 18, 30]. Using Markov State Modeling (MSM) techniques Plattner et. al. [17] were able to assess the formation of the Barnase-Barstar dimer. However, while very powerful, MSM techniques do not have direct access to the full dissociation transition due to the high barriers involved, which in turn induce long dwell times in the bound states. In contrast, the Transition Path Sampling (TPS) methodology bypasses the long dwell times in the stable unbound and bound states by focusing on the reactive association/dissociation trajectories directly [31]. TPS harvests a collection of unbiased molecular dynamics trajectories connecting two predefined stable states. The resulting path ensemble contains all pertinent dynamical and mechanistic information.

Here, we apply TPS to the rare dissociation/association transition (k_off_ ≤ 0.1 *s^−^*^1^) of the widely experimentally studied *β*-lactoglobulin (*β*-lac) globular protein dimer. We obtain atomistic insight in the mechanism of this transition by analysing the dynamically unbiased rare pathways and by extracting the best low dimensional models for the pertinent reaction coordinates (RC). This analysis includes the role of the solvent during the dissociation/association process itself, something that is only possible because we have access to the reactive transition paths. In addition, our study provides general insight in the dissociation mechanism for proteins that do not bind through steering interactions [6, 32].

We find that dissociation of the *β*-lac dimer from its native state occurs along a variety of multistep routes with transient intermediates, namely a direct *aligned* unbinding route and an indirect dissociation route through *sliding* or *hopping* to misaligned configurations before unbinding. The first dissociation bottleneck or transition state ensemble (TSE) appears in all paths and is associated with the breaking of native contacts and the formation of HB-bridging water mediated interactions. There is a secondary bottleneck related to the solvation of the persisting salt bridge R40-D33. In the *aligned* mechanism the number of waters at the interface and the distance r_40*−*33_ are pertinent ingredients of the RC for this mechanism. In the sliding mechanism the secondary bottleneck involves also a relative rotation of the proteins, which induces a loss of native contacts while still forming the R40-D33 directional salt bridge. Here, the distance r_40*−*33_ and the rotation angle *ϕ* are important ingredients of the RC. Finally, the hop mechanism involves a rebinding before dissociating, with the protein-protein distance as the relevant RC. As the TPS pathways are microscopically reversible we can interpret the transition paths both in the forward (dissociation) as well as in the backward (association) direction. However, we stress that these paths do not represent the complete, full association/ dissociation process, but only those that occur within a maximum allowed limited time, as specified by the path ensemble.

Indeed, the presence of multiple sequential barriers causes paths to become long, hampering the transition path sampling. To avoid the additional dwell time between the first barrier, the breaking of the contacts, and the secondary solvation barrier, we also sampled the transition path ensemble for this secondary barrier using a more relaxed non-specifically bound state definition (abbreviated B) using the desolvated (dry) contact area. The trajectories in this path ensemble are significantly shorter, but also exhibit mechanisms involving *direct* dissociation, a *sliding* mechanism where first the dry area decreases by a sliding movement, before dissociation, and a *hopping* dissociation route where the proteins first rebind before completely dissociating.

Since we find that solvent is an ingredient in the RC, e.g. the number of interfacial waters, or hydrogen bond bridging water occurring in the transition state ensemble, we further investigate the structure and reorientation dynamics of water during the dissociation process. Here, we find that water at the native dimer interface comprises a *disordered slow* population due to formation of long-lived hydrogen bonded bridging water, and a *tetrahedral fast* population, reorienting faster than bulk, which is typical of hydrophobic solvation and thus characteristic of *β*-lac’s mixed polar-apolar interface. This finding also extends to water populations in the transition state, which contain more mobile hydrogen bond bridging waters, enabling enhanced rotational mobility of the protein dimer with respect to a completely dry contact surface.

The remainder of the paper is organised as follows. In the following section we review the MD simulation settings and TPS algorithms as well as the analysis methods and tools. Next, we present and discuss the results on the specific unbinding transition and the non-specific binding transition. This is followed by a analysis of the hydration structure and dynamics. We end with concluding remarks.

## Methods

### Molecular Dynamics

In this study we performed molecular dynamics (MD) simulations using Gromacs 4.6.7 with GPUs [33]. All potential energy interactions were defined using the amber99sb-ildn and TIP3P force fields [34, 35]. We obtained the *β*-lactoglobulin (*β*-lac) PDB structure from the Protein Data Bank (PDB:2AKQ) and placed it in a dodecahedral simulation box which was energy minimized in vacuum using the conjugate gradient method. After solvation of the box with 20787 water molecules and a second energy minimization, we performed a 10 ps NPT short equilibration of water under ambient conditions with the protein position restrained. The solvated system was equilibrated for 1 ns in ambient conditions in the NPT ensemble and thereafter was subjected to a long 200 ns NPT simulation. All bonds were constrained using the Lincs algorithm. We used a 1 nm cutoff for the non-bonded Van Der Waals interactions. The electrostatic interactions were treated by the Particle Mesh Ewald algorithm, with a Fourier spacing of 0.12 nm and a 1 nm cutoff for the short range electrostatic interactions. The updating frequency of the neighbour list was 10 fs with a cutoff of 1 nm and the time step was 2 fs [34]. In the NPT simulations the temperature was controlled using the velocity-rescaling thermostat[36] with a coupling time constant of 0.2 ps. The pressure was controlled using the Parrinello-Rahman barostat[37] with a coupling time constant of 1.0 ps.

Short NVE MD simulaiton were performed to characterise the water structure and dynamics. No position restraints were imposed, and in order to prevent energy drift we used a switching function for the non-bonded interactions from 0.8 to 1.0 nm. The pair lists were updated every 5 fs with a cutoff of 1.2 nm, and the time step was 1 fs. The frequency of the energy calculation was 10 fs.

### The TPS spring shooting algorithm

Transition Path Sampling [38, 39] (TPS) harvests an ensemble of rare trajectories that lead over a high free energy barrier, connecting two predefined stable states. Starting from an initial reactive path, TPS performs a random walk in trajectory space by selecting a time frame, changing the momenta slightly and shooting off a new trial trajectory forward and backward in time by integrating the equations of motion. Acceptance or rejection of the trial trajectory is done according to the Metropolis rule [38, 39] which for the standard two way shooting move under fixed path length the trial move is accepted if the trial path connects the two stable states. If not the trial path is rejected.

The more efficient one-way flexible shooting algorithm [39, 40] samples the minimal length pathways between stable states and has been previously used in other protein systems [41, 42]. The one-way shooting method has several drawbacks. First, it requires more shots to decorrelate paths (although not more computer time). Second, it suffers in efficiency for asymmetric barriers, which occur, for instance, when the system on one side of the main barrier is trapped in an intermediate state, while it can easily reach the stable state on the other side. This means the paths on the trapped side become much longer. When uniform one-way shooting is used, this asymmetry leads to many more shooting attempts on one side of the barrier with respect to the other, increasing the inefficiency.

The spring shooting algorithm is especially developed for use with the one-way algorithm[43]. It only differs in the way the shooting point is selected. Instead of uniform random selection, the spring shooting shifts the shooting point index with respect to the last successful shooting point, not in a symmetric but in a asymmetric way according an acceptance criterion

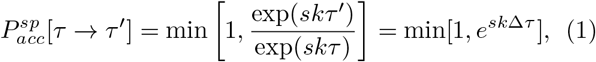

where ∆*τ* = *τ′* – *τ* is the number of shifted frames from the previous shooting point *τ*, *k* denotes a force constant determining the magnitude of the bias, and *s* ∈ {–1, 1} is determined by the direction of shooting i.e. *s* = –1 for forward shooting, and *s* = 1 for backward shooting. The spring shooting algorithm thus treats the forward and backward shooting move as different types of moves. As a large ∆*τ* either yields an exponentially small acceptance ratio or is likely to produce a failed shot, in practice, we limit the choice of ∆*τ* between the interval [–∆*τ_max_*, ∆*τ_max_*], analogous to the maximum allowed displacement in a regular MC translational move. When the trial shooting point falls outside the current path the acceptance probability becomes zero, and the move is rejected. The remainder of the shooting move is identical to the uniform one-way shooting algorithm. For a detailed description of the algorithm see SI and Ref. [43].

The advantage of this approach is that unfavourable shooting points are discarded without extra cost. Pathways are decorrelated as much as possible, without wasting time creating partial paths that do not contribute to the decorrelation. Note also that the algorithm rejects trial paths which become longer than *L_max_*, which is set to prevent memory or storage problems, or as an indication that the path generation went awry, e.g. became trapped in an long-lived intermediate state.

### Defining the stable states and creating the initial path

To define the stable states we performed 200 ns MD in the NPT ensemble at ambient conditions, during which *β*-lac dimer remained in its native bound state (see Figure. S1). The native contacts were identified as those residue pairs that stayed within a minimum heavy atom distance of 0.4 nm for at least 90% in the 200 ns NPT trajectory. As shown in Table S1, only 8 residue pairs are shown to fulfill this criterion (150-146, 148-148,146- 150,148-147,147-148,149-146,146-149,33-33). These contacts are between residues of the beta sheets of the I-strand, and the AB-loops of the protein and have been also verified to be important for the stability of the dimer by experiments [29]. These eight residue pairs, as well as four native hydrogen bonds (between backbone NH and CO of residues 146-150,148-148,150-146) define the stable native contact state (N) Note that residue pairs 33-40 and 40-33 discussed in the Results section have smaller occupancy than 70%, and hence are not part of the native state. The unbound state (U) requires the minimum distance between the two proteins to be greater than 1 nm (*r_min_ >* 1 *nm*). Finally, the non-specific dissociation 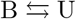 transition requires the definition of a bound state B. To be as non-specific as possible, we only required for this bound state that the protein-protein interfacial area is *>* 2 (nm^2^). All definitions are summarized in Table 1).

**TABLE I:** State definitions for the native state (N), the unbound state (U) and any bound state (B) as a function of native contacts (NC), native hydrogen bonds (N H-bonds), protein-protein minimum distance (*r_min_*) and dry contact area.

| | NC | N H-bonds | $r_{min}$ | Dry area |
| --- | --- | --- | --- | --- |
| Native (N) | $\geq 8$ | $\geq 4$ | - | - |
| Unbound (U) | 0 | 0 | $> 1 \text{ nm}$ | 0 |
| Bound (B) | - | - | - | $2 \text{ nm}^2$ |

To obtain the initial path for TPS we enforced the dissociation from the native bound state using Metadynamics at 300 K employing the PLUMED package[44] with the above-mentioned MD settings. As the resulting trajectory is strongly biased, we launched unbiased MD trajectories from particular frames, performed at a slightly elevated temperature of 330 K to avoid getting trapped in long-lived near-native intermediate states. Concatenating a trajectory returning to the native state with one going forward to the unbound state yields the desired unbiased initial TPS path. For completeness we checked that performing the simulations at 330 K does not perturb the protein conformations significantly. See the SI for more details.

### TPS simulation settings

We performed TPS simulations of the native state to the unbound state 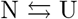 at *T* = 330 K, *P* = 1 atm in the NPT ensemble using home written scripts encoding the spring shooting scheme. The maximum path length for this transition was set to *L_max_*=7000, which, with frames saved every 10 ps, translates into a maximum path duration of *t_max_* = 70 ns. The spring shooting move parameters were set to *k* = 5 and ∆*τ_max_*=200. The spring constant ensures the shooting points remain close to the top of the very asymmetric barrier of *β*-lac dissociation. The ∆*τ_max_* was chosen small compared to the maximum path length allowed (3%) so that the shooting point rejection as well as the whole trial move rejection was kept to a minimum. For the 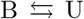 transition, the spring constant *k* is 0.1 and the ∆*τ_max_* is 70 frames.

### Analysis of the path ensemble

Home-written scripts analyzed the path sampling results to produce the path tree, the least changed path (LCP), the path length distribution, and the path density. We construct the path density by choosing two order parameters (e.g. protein-protein minimum distance vs native patch vector angle) and binning each frame of each trajectory in the path ensemble to a 2D grid. Every path can only contribute to a specific bin once, even if visited multiple times. Note that accepted paths can occur multiple times in the ensemble, depending on whether the next trial moves have been rejected. The least changed path (LCP), consisting of the stretches between successive alternating forward/backward shooting points acts as an approximation for the transition state ensemble[43]. For the path length distribution each accepted path of different length *L* is histogrammed according to its weight in the path ensemble.

Since the protein orientation degrees of freedom might be important during the dissociation transition/association, we calculate the relative orientation of the two protein, characterized by an angle *ϕ* (see Figure S7 for a graphical illustration)

Water plays an important role in dissociation/association. In order to address the solvent degrees of freedom, for each configuration we count the number of waters residing inside a cylindrical tube between the two proteins. The tube’s base centers are defined by the center of mass of each protein, a radius *r* = 1.4 nm, or *r*=1.1 nm and length *L* being the centre of mass distance between the two proteins.

### Reaction coordinate analysis by Likelihood Maximization

The reaction coordinate is a invaluable description of a complex transition as it can predict the progress of the reaction. Transition Path Theory (TPT) states that the perfect reaction coordinate is the commitment probability (committor or p-fold) *p_B_*(*x*) as it gives the probability for a configuration *x* to reach the final state B [45]. Although the committor *p_B_*(*x*) is mathematically well defined, it is excessively expensive to compute, and being a high dimensional function, not very insightful as to which are the relevant slow degrees of freedom. Peters and Trout developed a Likelihood Maximization (LM) method to extract the best (linear) model for the reaction coordinate based on an approximation of the committor function using the shooting point data from TPS[46, 47]. Each trial shooting point in this data set can be regarded as drawn from the committor distribution [46, 47]. Using as input the N forward (or backward) shooting point configurations *x_sp_* of the accepted trajectories ending in the final state B (*x_sp_ → B*) and the shooting points of the rejected trajectories ending in state A (*x_sp_ → A*), the method defines the likelihood that a model reaction coordinate *r* can reproduce the observed data

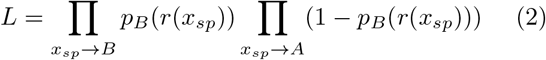

where the committor *p_B_*(*r*) is a function of the reaction coordinate *r*. The reaction coordinate *r* is modeled/parametrized as

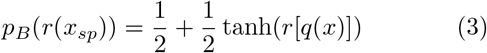

where the reaction coordinate *r*(*q*(*x*)) is approximated by a linear combination of *m* collective variables *q*(*x*) as follows.

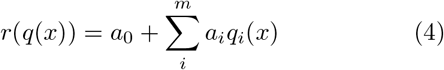

The LM analysis serves as a screening tool for linear combinations of candidate CVs and returns the one that best parametrizes the committor probability, given a good dataset obtained from the TPE. Adding an additional CV in the analysis, i.e. increasing *m* by one, should lead at least to an increase of 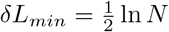 in the maximum likelihood, in order for the RC to be deemed a significant improvement [46, 47]. The spring shooting algorithm is naturally suited for use with this LM approach, since it gives access to shooting points close to the transition state ensemble. We use the candidate CVs listed in Table S2.

### Water structure and dynamics analysis

To obtain insight in the role of the solvent we analyzed the structural and orientation dynamics of water in the dissociation transition, with the same protocol as in Refs. [48, 49].

As the TPS simulations are done at 330 K, we first analyze the water structure and dynamics at this temperature as follows. From a decorrelated reactive path in TPS ensemble we selected three different frames belonging to the native, the near native transition state TSR_2_ (Native contacts=2, *ϕ*=60°), and the unbound state, respectively. For each of them a 1 ns NPT run at 330 K is performed with the proteins position restrained, followed positions of the short 1ns NPT simulations (see MD section for details). Frames were saved every 100 fs in order to obtain sufficient data for analysis of the water dynamics. In order to identify whether water structure and dynamics changes significantly from 330 K to 300K, we repeated the same analysis at 300 K.

From the short NVE trajectories we computed the re-orientation decay time *τ* for each water molecule in the hydration layer of the proteins from an orientational correclation function (see SI for details). The decay times of the individual water molecules allow us to establish the relation between the water structure and dynamics.

In addition we characterize the tetrahedral structure of water around amino acids based on the probability distribution *P*(*θ*) of the minimum water-water OOH angle *θ* (see Figure. S8) for all water-water pairs within 3.5 *°A* from each other and solvating the amino acid [50–52]. The distribution *P*(*θ*) of these angles takes on a bimodal distribution with a minimum at 30°, distinguishing between tetrahedral water population (angles lower than 30°) and a perturbed H-bond network, mostly occurring around hydrophilic groups (angles higher than 30^*°*^). The tetrahedral structure parameter *S* is defined as the integral of *P* (*θ*) up to *θ* = 30° [50–52]. Water around hydrophobic groups has a larger *S* due to smaller H-bond angles *θ*, inducing stronger water-water bonds, with larger energy fluctuations and therefore a positive heat capacity of the solvating water. In contrast, the introduction of a hydrophilic group around water strains the water-water H-bond angle and shifts the angle distribution to higher values, and hence a lower *S*, thus decreasing the water-water bond energy and fluctuations which decreases the heat capacity of solvation [50–52]. Throughout the text we will associate tetrahedral structured water with a large *S* value (high tetrahedral water population) and unstructured water with a low *S* value (low tetrahedral water population). Unstructured water coinciding with slow reorientation dynamics (as characterized by *τ >* 4 ps) will be labelled as *disordered* water.

We computed the distribution *P*(*θ*) and extracted a structural order parameter *S* for each molecule hydrating a amino acid separately. We bin the *τ – S* pair for each residue in a 2D histogram, in order to investigate possible correlation between water tetrahedral structuring and reorientation dynamics (see SI for more information).

### Hydrogen bond bridge survival correlation function

The hydrogen bond bridge correlation function (eq. 5) is a correlation function that traces the decay time of a hydrogen bond bridge between two intermolecular protein residues.

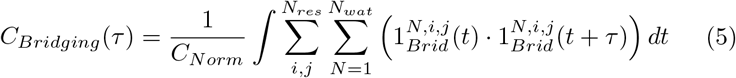

Where *τ* is time, *C_Norm_* is a normalisation constant, such that *C*_Bridging_(0) =1, *i* and *j* are running over the residue number of proteins *A* and *B* respectively, *N_res_* is the total number of residues per protein and *N_wat_* is the total amount of waters in the simulation box. 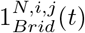 is an indicator function at time t, which is unity if water *N* is hydrogen bond bridging residues *i* and *j* and zero otherwise.

## Results and Discussion

### TPS of the specific dissociation transition

We performed several TPS runs at 330 K employing the spring shooting algorithm for the transition between the unbound state (U) and the native bound dimer state (N). In total we performed 560 shooting trial moves, of which 18.3 % was accepted with total aggregate and accepted path simulation time of 23.7 and 3.43 *μ*s respectively. Decorrelation was tested using path trees (See SI). The average path duration was 33.6 ns. The path length distribution in Figure. 1 is broad and includes a significant population of longer paths. To shed light on this bimodal distribution, we analyse the transition path ensemble (TPE), first by inspection.

**FIG. 1:**
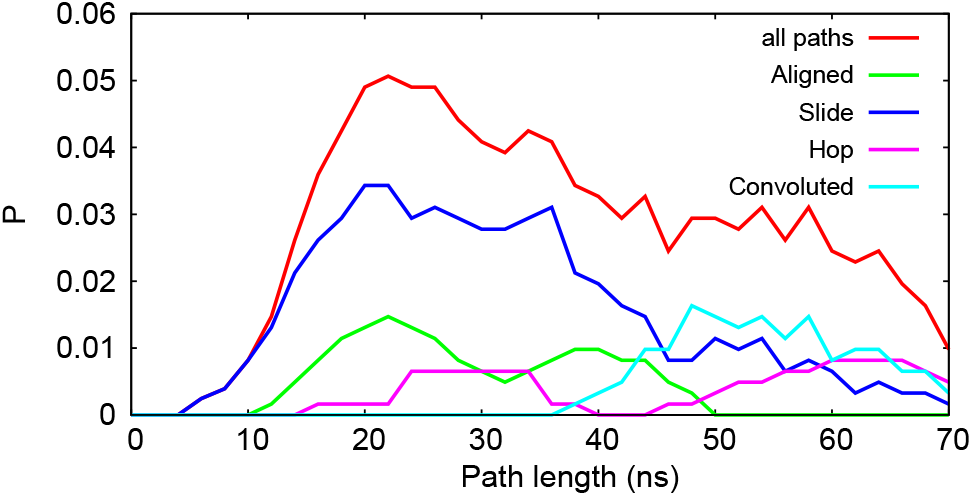
Path length distribution of the 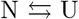 path ensemble and for the different observed mechanisms.

The trajectories in the TPE all fall into three categories, representing three qualitatively distinct dissociation mechanisms. 1) In the *aligned* mechanism proteins directly separate to the unbound state without (much) rotation. 2) In the *hopping* mechanism proteins first dissociate and then rebind to a low dry surface non-specific configuration before fully dissociating. 3) In the *sliding* mechanism proteins first rotate and slide out of the native state to a higher dry interface non-specifically bound state, henceforth called transient near native state, before fully dissociating. Figure. 2 illustrates these three mechanisms schematically. Switches between these different mechanisms frequently occur in the path sampling (see Table S3). We also observe trajectories exhibiting a convolution of the above mechanisms, in which the proteins slide and rotate away from the native state, but instead of dissociating, return to the native state, followed by an aligned dissociation. The sliding mechanism is most abundant in the TPE. The aligned mechanism is least prevalent, most likely due to the lower orientation entropy. The longest transition paths tend to be the sliding, hopping and convoluted mechanisms, because of the presence of transiently formed intermediates. While sliding and rebinding binding mechanisms have been reported previously [17, 19, 25, 26], here we observe all three mechanisms simultaneously. To gain further insight we define several collective variables (CVs) that are important for the dissociation mechanisms. Besides the number of native contacts, NC, and the minimum distance *r*_min_, we define the angle of rotation *ϕ* where *ϕ* = 0° corresponds to the fully aligned dimer. Further, we identify specific contacts. Many trajectories in the path ensembles exhibit the transient but relatively long-lived (occupancy *>* 10 ns and heavy atom distance *<* 0.4 nm) R40-D33 double salt bridge contact between the carbonyl groups of D33 of one protein and the amide groups of R40 of the other protein. Other long-lived intermolecular contacts are preserved in a large number of paths, although not throughout the entire TPE (see Table 2). Note the lack of symmetry due to imperfect sampling. We highlight a configuration containing the three most occurring contacts (R40-D33, H146-S150, I29-S150) in the path ensembles in Figure. 3. These findings signify that the dissociation/association process is not a one step process, and transiently formed interactions occur during the pathway, something that is also observed for the Barnase-Barstar dimer, the Insulin-dimer, and Ras-Raf-RBD, RNase-Hi-SSB-Ct, TYK2-pseudokinase complexes [17, 19]. Sakurai et. al. [29] showed that the R40- D33 interaction is important for the dimer formation process, as mutating any of these residues to an oppositely charged amino-acid drastically reduced the binding constant. Also native contact residues H146, R148 and S150 are important for dimerization, as the binding constant decreases when these residues are mutated to proline, thus breaking the *β*-sheet structure at the native binding site. Indeed we observe these contacts as well (see Table 2 and Table S1). As interactions of these residues H146, R148 and S150 are included in the definition of the native contacts, we focus here on the salt bridge distance *r_R_*_40*−D*33_. (Note that we could as well have focused on the symmetric salt-bridge D33-R40).

**FIG. 2:**
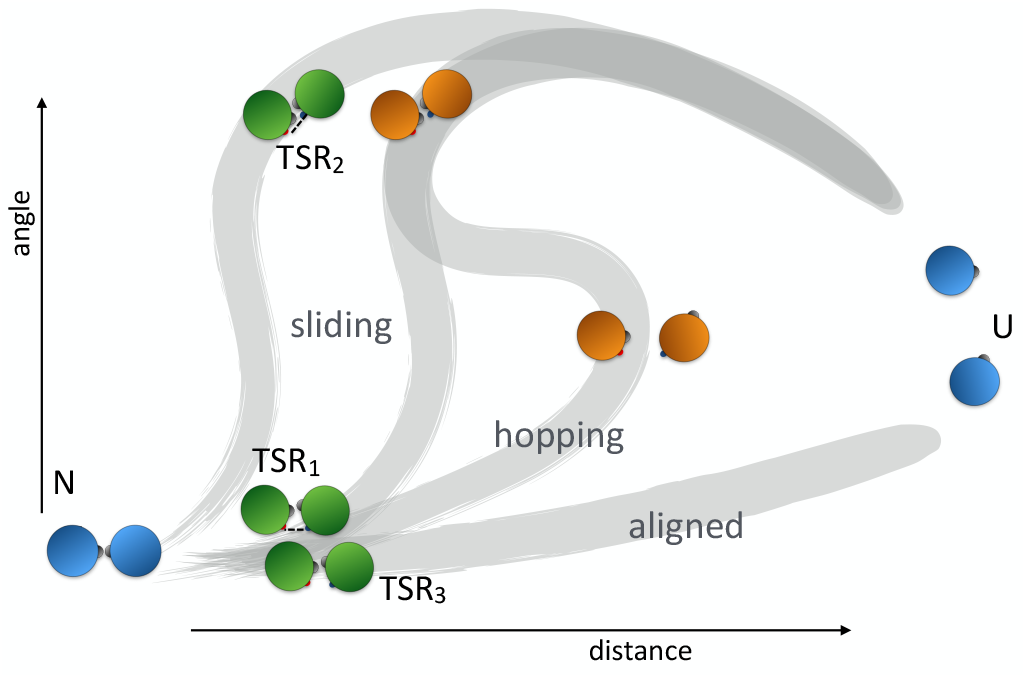
Cartoon network of transitions and respective TSRs during the full dissociation/association process. We identify three types of mechanisms: the aligned, hopping and sliding transitions.

**FIG. 3:**
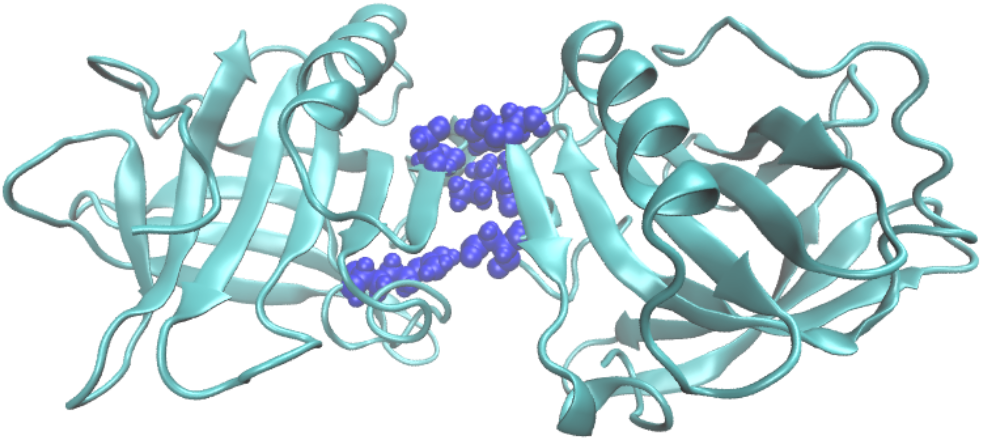
Structure of an on pathway transiently formed intermediate. In blue are highlighted the long-lived contacts H146-S150, R40-D33, I29-S150.

Since the role of water in hydrophilic association [10, 23, 24] is crucial, yet elusive, we define several solvent based CVs: *A*_dry_, the dry surface area of contact between the proteins, *N_HB_*, the number of hydrogen bonded bridging waters, and 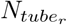, the number of waters in a tube of a certain radius *r* between the proteins centres of mass. Figure. 4 shows the path density as a function of a variety of CVs for both the full transition path ensemble (TPE) as well as for the least changed path ensemble (LCP) (see Methods and SI). The LCP approximately samples the transition barrier region, serving as a proxy of the transition state ensemble [43]. The path densities of the full TPE show that all paths pass through configurations with partially formed native contacts (1*< N C <*4), but the LCP indicates that in fact the TSE is split into several regions, which we denote transition state regions (TSR). TSR_1_ occurs at *ϕ <* 40° and 1 *< N C <* 4, while TSR_2_ is located at *ϕ >* 50° and 0 *< N C <* 2. At the same time the TSRs are characterised by a substantial number of hydrogen bonded bridging waters. While this CV is not found to be among the most pertinent ingredients of the RC discussed later, the formation/breaking of hydrogen bond bridging waters between the proteins represents a dynamical bottleneck in the association/dissociation process. This observation is in agreement with the prediction of Ben-Naim and Northrup [23, 24] that water-mediated interactions drive or characterise the hydrophilic association. The path density and LCP in the *ϕ − r*_40*−*33_ plane, suggest the presence of a third TSR_3_ characterised by a hydrogen bonded water between residues R40 and D33. Several representative configurations from these TSRs are highlighted in Figure. 4. The TSR_3_ bottleneck suggest that water solvation of R40-D33 helps the proteins escaping a very strong and directional salt bridge interaction during the dissociation process.

**FIG. 4:**
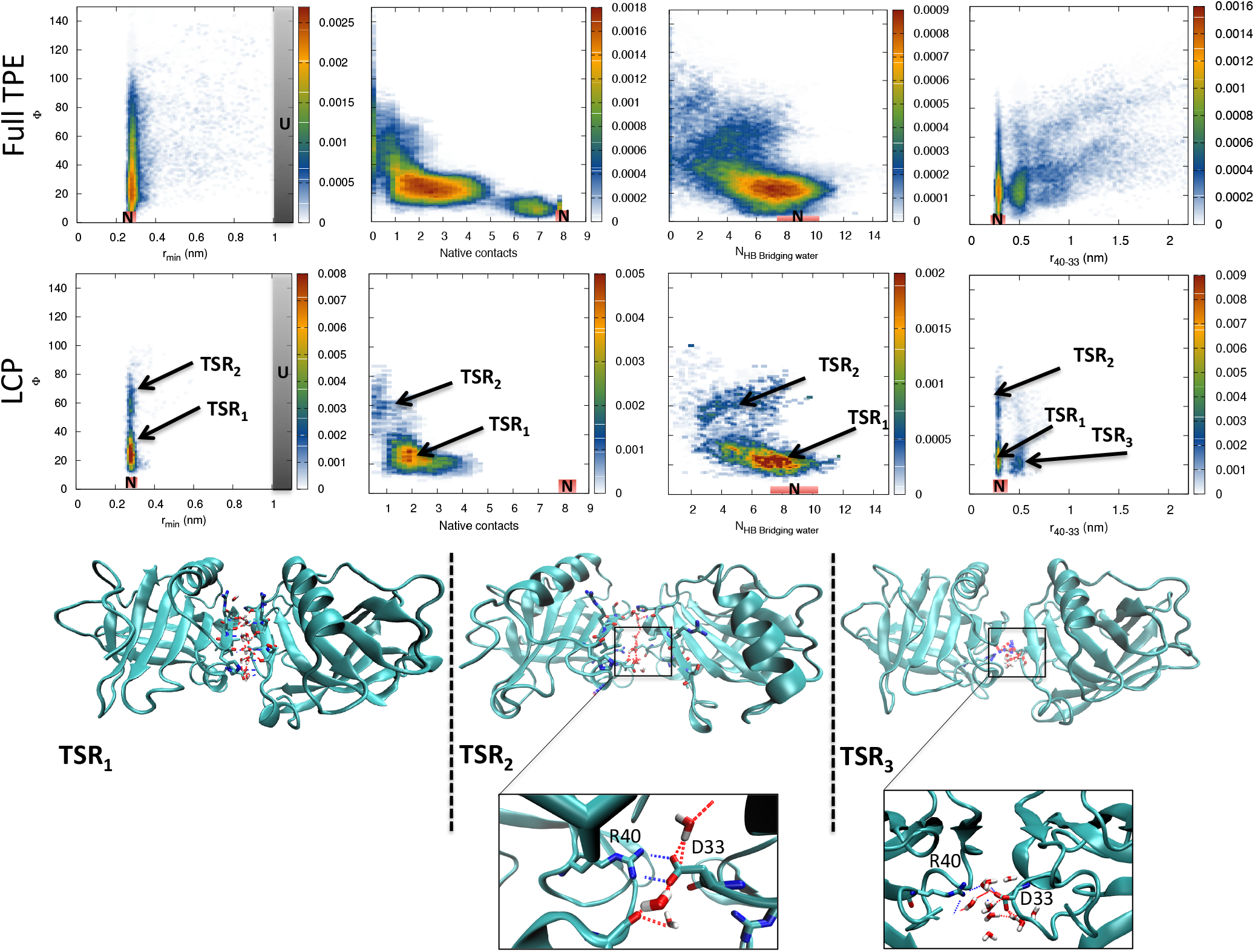
Path density plots of *ϕ* as a function of protein-protein minimum distance (col 1), native contacts (col2) number of hydrogen bond bridging waters (col3) and R40-D33 distance (col4) for the full transition path ensemble (full TPE) and least changed path ensemble (LCP) respectively. The third shows snapshots of configurations of the TSR1, TSR2 and TSR3.

**TABLE II:** Number of paths in which individual intermolecular contacts between aminoacid of protein A with aminoacid of protein B occur with a lifetime higher than 10 ns, for each path ensemble.

| #aa<sub>A</sub>-#aa<sub>B</sub> | TPE | #aa<sub>A</sub>-#aa<sub>B</sub> | TPE | #aa<sub>A</sub>-#aa<sub>B</sub> | TPE |
| --- | --- | --- | --- | --- | --- |
| H146-S150 | 77 | D33-R40 | 47 | I29-F151 | 32 |
| R40-D33 | 71 | S150-H146 | 44 | S150-I29 | 28 |
| I29-S150 | 67 | I29-Q155 | 44 | H146-N152 | 24 |
| D28-N152 | 53 | Q155-I29 | 35 | K141-D130 | 22 |

As the specific dissociation TPE is a superposition of the three possible mechanisms, separately plotting the TPE for each route in Figure. S11 enables a discussion of each route individually. For each of these sub-ensembles we also performed a reaction coordinate analysis (see Methods and SI). In the aligned mechanism the native contacts break, while the angle *ϕ* stays below (*ϕ <* 50°), followed by solvation of the dry contact surface and unbinding of the proteins. The paths pass through TSR_1_ where the proteins have partially formed native contacts (1 *< N C <* 5) and through TSR_3_, which involves the breaking of the salt bridge between R40 and D33, and forming a water mediated interaction. The dynamical bottleneck upon binding is thus the correct alignment, local rearrangement and formation of the native contacts, as well as expelling the water at the interface, and in particular at the R40-D33 contact, and increasing the number of hydrogen bond bridging water. RC analysis shows the solvation of the protein-protein interface (N_*tube*11_) is the most important RC for the aligned mechanism (see Table 3 and Figure. 5). However, the salt bridge distance *r_R_*_40*−D*33_ is deemed a good additional reaction order parameter, only barely missing the threshold for significance, perhaps due to the presence of TSR_3_ (see Table 3). In the hopping mechanism the dimer first breaks the native contacts to become dissociated with zero dry contact surface area, but with several hydrogen bond bridging waters. Then the proteins rebind to a non-specific low dry interface configuration characterised by a large angle *ϕ >* 50°, a small dry contact surface area 1 − 2 nm^2^, and some hydrogen bond bridging waters. Finally the dimer completely unbinds. The paths pass through TSR_1_ and TSR_3_ as proteins dissociate and rebind. RC analysis for the hopping mechanism indicates that the protein-protein centre of mass distance is the most pertinent collective variable (see Table 3). In the sliding mechanism proteins first break their native contacts by sliding and rotating to a misaligned configuration before unbinding. During sliding a dry contact surface area is preserved (2-5 nm^2^) with some hydrogen bond bridging waters present. Paths can pass through either TSR_1_, TSR_2_ and TSR_3_ with roughly equal probabilities. The R40-D33 salt bridge is still present in TSR_2_. RC analysis indicate the relevant CV describing the bottleneck is the salt bridge distance as well as the rotation angle *ϕ*. Thus, either water can assist breaking of directional interactions-(R40-D33) present in TSR_1_ before rotating, or vice versa in TSR_2_.

**FIG. 5:**
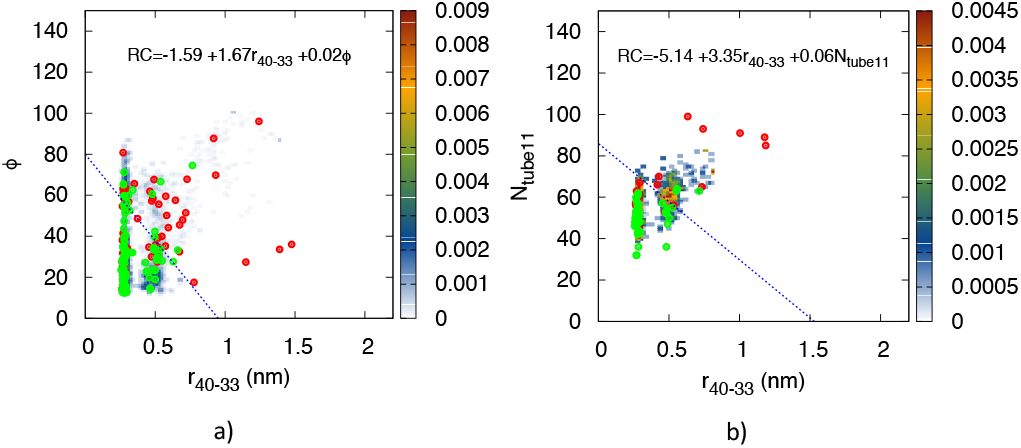
Plot of forward shooting points, and predicted dividing surface *RC* = 0 (black solid lines) for a) the sliding and b) aligned paths. Red points end in U, green points in N. Note the split nature of the TSE.

**TABLE III:** LM analysis for the 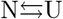 transition based on the forward shooting points for a) the aligned paths b) the sliding paths and c) the hopping paths. *δL_min_* denotes the minimum required increase in likelihood when adding an additional CV.

| n | ln L | reaction coordinate |
| --- | --- | --- |
| Aligned paths $\delta L_{min} = 2.04$ (59 shooting points) | | |
| 1 | -33.24 | -5.24 + 0.09N<sub>tube11</sub> |
| 2 | -31.46 | -5.14 + 3.35r<sub>40-33</sub> + 0.06N<sub>tube11</sub> |
| 3 | -31.32 | 23.24 - 9.8 Dist + 2.95r<sub>40-33</sub> + 0.14N<sub>tube11</sub> |
| Sliding paths $\delta L_{min} = 2.54$ (161 shooting points) | | |
| 1 | -104.93 | -3.41 + 0.02N<sub>tube14</sub> |
| 2 | -102.28 | -1.59 + 1.67r<sub>40-33</sub> + 0.02ϕ |
| 3 | -102.09 | -2.44 + 1.84r<sub>40-33</sub> + 0.27Native contacts + 0.03ϕ |
| Hop paths $\delta L_{min} = 1.74$ (33 shooting points) | | |
| 1 | -18.95 | -15.17 + 4.30Dist |
| 2 | -17.83 | -35.21 + 12.61Dist - 0.05N<sub>tube14</sub> |
| 3 | -17.25 | -78.05 + 19.44Dist + 1.98Native contacts + 27.50r<sub>min</sub> |

### TPS of the non-specific dissociation transition

In the previous section we focused on the specific dissociation/association from and to the native dimer state. In this section we focus on the non specific dissociation/association of this dimer by studying the 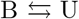 transition. Here the non-specific bound state B is defined in a much less strict sense by requiring a minimal de-wetted dry contact surface area of 2 nm^2^ (see Table 1). We performed a TPS run at 300 K employing the spring shooting algorithm for the non-specific dissociation/association mechanism, between the unbound state (U) and any bound state (B). In total we performed 531 shooting trial moves, of which 37 % was accepted, with total aggregate and accepted path simulation time of 854 ns and 398 ns respectively. The path length distribution in Figure. 6 shows that the average path length is now 4.3 ns, an order of magnitude shorter than in the 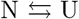 transition. A partial path tree is shown in Figure. S12. Upon inspection the path ensemble shows three distinct mechanisms that are very similar to the 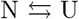 case, namely a *direct* dissociation mechanism, one that involves *hopping* and rebinding mechanism, and the *sliding* mechanism, depicted in a cartoon representation in Figure. 7. In the *direct* mechanism proteins transit in around a 1 ns between a unbound and bound state, by quickly increasing the dry contact surface area. In the *hopping* mechanism proteins solvate the dry contact surface area, (partially) unbind and rebind first before fully unbinding. The hopping paths are not only the most abundant, they tend to be the longest, because they undergo a two step process. The reason the hopping transition is more abundant than in the 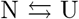 transition, is that the 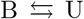 transition has a much less strict bound state, which can be easily reached via hopping. In the *sliding* mechanism a bound proteins first slide to a configurations with low dry contact surface area before dissociating.

**FIG. 6:**
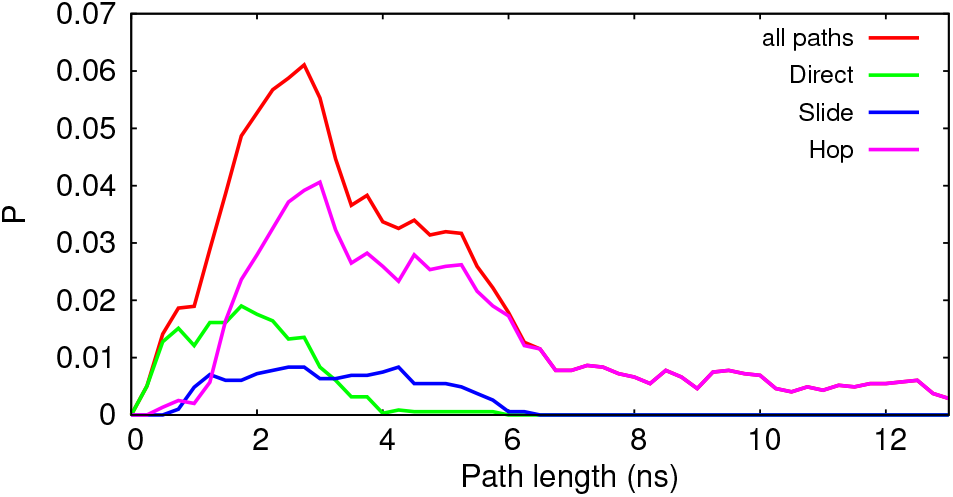
Path length distribution of the 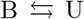 transition path ensemble with respect to the different underlying mechanisms. In red are all paths of the path ensemble, in green, blue, magenta, the ones corresponding to the direct, slide, hop dissociation/association.

**FIG. 7:**
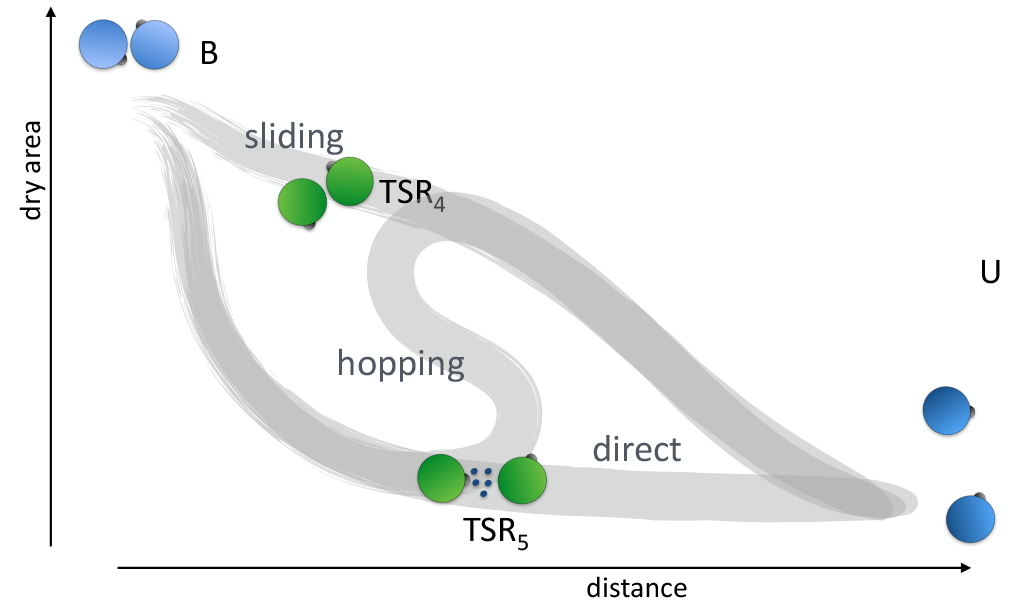
Cartoon network of transitions and respective TSRs during the non specific dissociation/association process. We identify three types of mechanisms: a direct, hopping, and sliding mechanism.

To analyse the TPE of the non-specific unbinding process 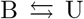, we computed several collective variables. Besides the minimum distance *r_min_*, the angle of rotation *ϕ*, we measure the minimum salt bridge distance, the dry contact surface area *A_dry_*, the number of waters in a cylinder of a certain radius *r* between the proteins centres of mass, *N* tube_r_ and the number of hydrogen bond bridging water *N*_HBbridgingwater_. Figure. 8 shows the path density for both the LCP and for the entire transition path ensemble for several combinations of these CVs. As these path densities are convolutions of the three different mechanism we also present multiple TPE path density plots for each individual mechanism in Figure. S13. The LCP shows two major TSRs, one (TSR_4_) at r_*min*_=0.3 and a dry area around 1-1.5 *nm*^2^, and one (TSR_5_) at r_*min*_=0.5 and dry area of 0.15 *nm*^2^. TSR_5_, is partly due to direct paths which show a simultaneous decrease in dry surface area, increased solvation, increased salt bridge distance upon dissociation and drastic decrease of the hydrogen bond bridging water (see Figure. S13). RC analysis of the direct mechanism (see Table 4) indicates that it involves formation of salt bridges and (to a lesser extent) attaining the proper orientation *ϕ*. As the direct mechanism only has to pass the barrier TSR_5_, this mechanism exhibits much shorter transition paths than the hopping transition (see Figure. 6). Visual inspection of configurations in TSR_5_ (see Figure. 9) indicated that while the two proteins are separated by a solvent layer without any contacts formed, their charged and polar residue dominated interface surfaces are correctly aligned, suggesting that from an association perspective the proteins can quickly form a dry area of more than 2 nm^2^. The path density for the sliding mechanism exhibits a strong peak at contact *r_min_* = 0.3 and dry area around 1-1.5 *nm*^2^, characteristic for TSR_4_. RC analysis of the sliding mechanism (see Table 4) shows that the pertinent degree of freedom is indeed the dry surface area. Moreover, this dry area involves formation of salt bridges as shown by the r_*min*_(salt-bridge) vs dry area plot of Figure. S13c. Figure. 9 suggests the TSR_4_ comprises formation of a few salt bridges in a wet interface without any other contacts formed, inducing a small crevice region around it. Hence the proteins have to sequentially slide each other in order to reach the bound state B. Hopping paths can exhibit both TSR_4_ and TSR_5_, as is clear by the presence of two peaks in the path densities of Figure. S13. Thus, the hopping transition can involve a barrier at contact, as well as a barrier at separated distances.

**FIG. 8:**
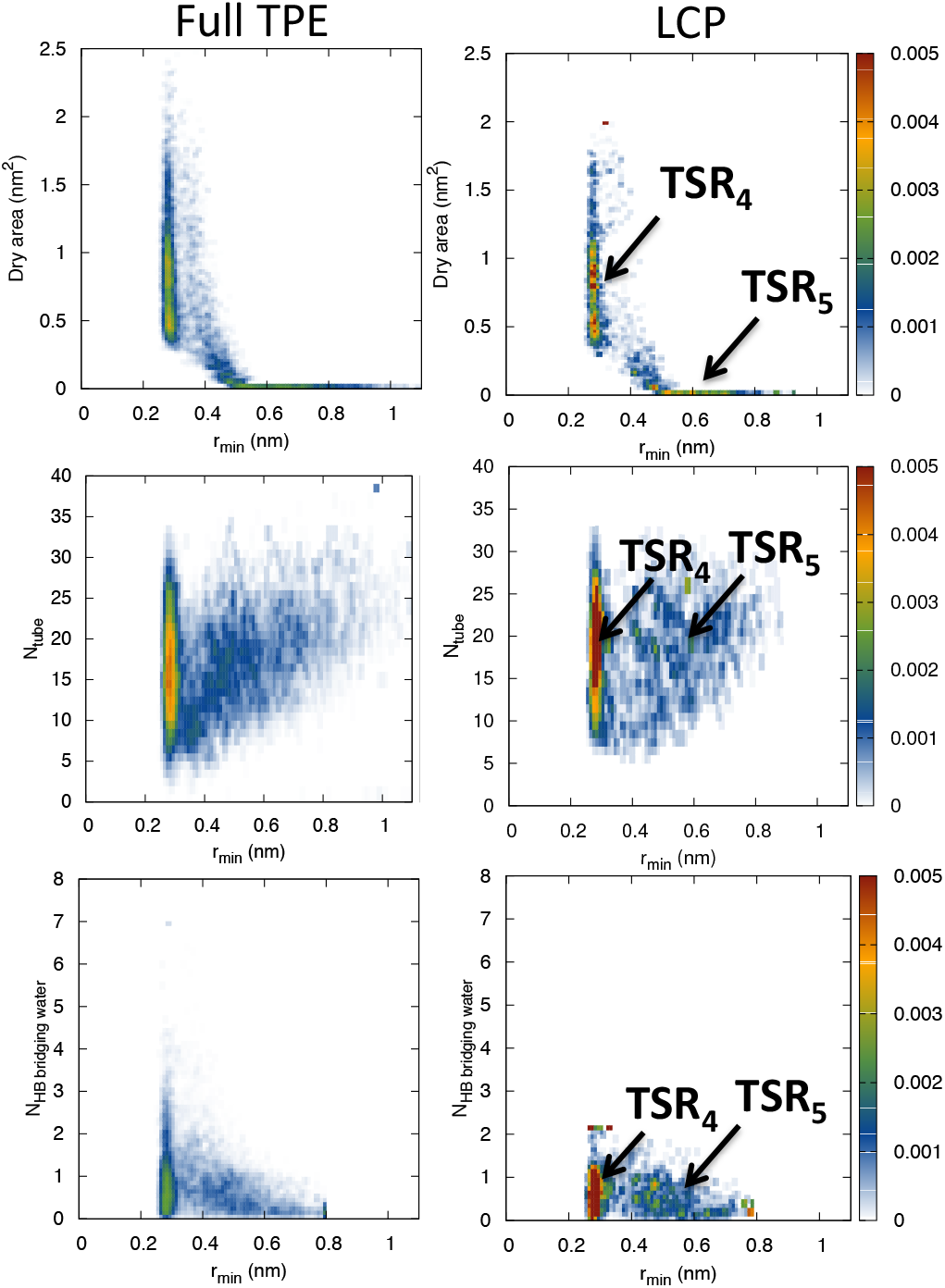
Transition Path Ensemble and Least Changed Path density plots of the number of dry area vs protein-protein minimum distance (up), number of waters in the tube of radius 0.8 nm vs protein-protein minimum distance (middle) and the Number of hydrogen bond bridging waters vs protein-protein minimum distance(bottom) for the TPE (left) and the LCP (right).

**FIG. 9:**
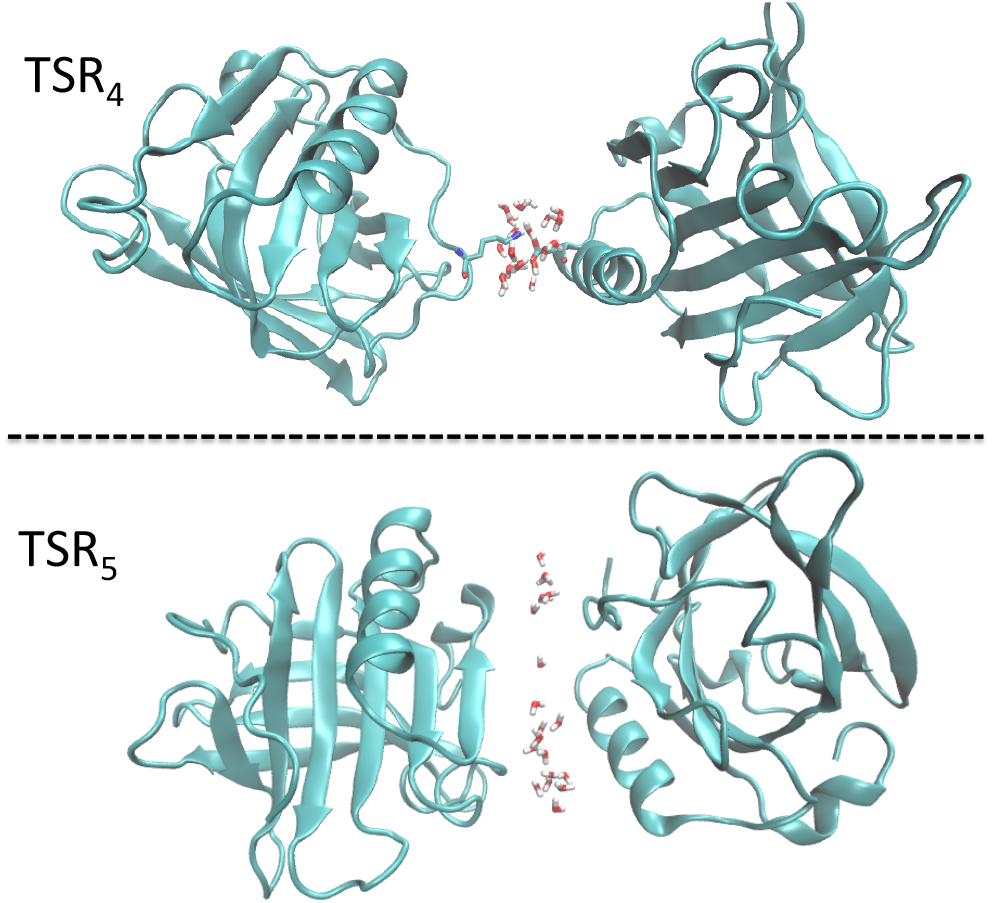
Snapshots of configurations of the TSR_4_ and TSR_5_.

**TABLE IV:** LM analysis for the 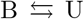 based on the Forward shooting points for a) the direct paths b) the sliding paths and c) the hop paths.

| n | $\ln L$ | reaction coordinate |
| --- | --- | --- |
| Direct paths $\delta L_{\min} = 2.07$ (63 shooting points) | | |
| 1 | -39.39 | 0.49 -0.24N |
| red!50 2 | -36.87 | 0.95 +2.85$r_{\min}$ (salt bridges) -0.10ϕ |
| 3 | -36.47 | 0.98 -0.15N +2.10$r_{\min}$ (salt bridges) -0.07ϕ |
| Sliding paths $\delta L_{\min} = 1.56$ (23 shooting points)) | | |
| red!50 1 | -14.00 | 0.93 -1.63 Dry area |
| 2 | -12.96 | 4.46 -4.64 Dry area -5.80$r_{\min}$ |
| 3 | -12.38 | 17.11 -1.02N -11.33$r_{\min}$ (salt bridges) +0.22ϕ |
| Hop paths $\delta L_{\min} = 2.55$ (164 shooting points) | | |
| red!50 1 | -108.42 | -8.50 +1.97Dist |
| 2 | -106.85 | -6.90 -0.78 Dry HF<sub>area</sub> +1.66Dist |
| 3 | -105.86 | -12.91 -0.89Dry HF<sub>area</sub> +1.93Dist +9.61S14 |

The difference between hopping and sliding is even more clearly visible in Figure. S14, which shows the LCP plot for each of these mechanisms. Indeed, while the sliding LCP shows basically just one peak at *r_min_* = 0.3 and dry area around 1-1.5 *nm*^2^, the hopping LCP exhibits a second peak at *r_min_* ≈ 0.6 and zero dry area. RC analysis of the hopping transition shows that the CM distance of the proteins is the most relevant collective variable (see Table 4). While this makes sense as hopping requires complete separation before rebinding, it is probably due to the mixed influence of the two TSRs. Again, we can view the sliding mechanism as rebinding without the proteins truly separating. Finally, we looked at the hydrophilic nature of the dry contact area, which, although it did not make the treshold also deemed a reasonable RC for the hopping mechanics. Plotting the LCP as function of dry interfacial area and the hydrophilic minus hydrophobic dry interfacial area in Figure. S14, it is clear that the nature of the dry interface is indeed hydrophilic, which is expected for a hydrophilic dimer former such as *β*-lac.

Note that hopping paths are in the majority in the path ensemble for the non-specific B-U transition (see Figure. 6), while the 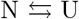 path ensemble showed more sliding paths (see Figure 1). We can rationalise this as follows. In the 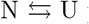 transition paths have to end in the N state, which causes an entropic bottleneck for the direct binding, and the hopping mechanism. Sliding requires a two dimensional search over the surface of the protein, which does not suffer from this entropic bottleneck as much. Hence, the sliding paths are the majority. In the 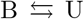 transition the bound state is not specific, and can easily be reached by the direct, sliding and hopping mechanisms. Hopping is now very likely as from a hydrated transition state the proteins can easily rebind. Our findings of direct, sliding and rebinding paths for the non-specific dissociation for the 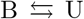, suggests that these atomistic mechanisms of protein unbinding, also occurring in the specific 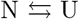 are general dissociation mechanisms. Indeed recent numerical work highlights these mechanisms using simplified models of protein binding[25, 26]. These findings are also fully compatible with previous MD simulations[6, 53].

### Structure and dynamics of hydration water during dissociation

As we find that water is an ingredient in the RC for both the 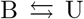 transition (for example in desolvation barrier TSR_5_), as well as in the 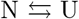 transition, and by the presence of hydrogen bond bridging contacts between proteins both at TSR_1_, TSR_2_ and TSR_3_, we further investigate the structure and reorientation dynamics of water during the dissociation process. We focus on the hydration of three states during the 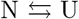 transition: a native, a transient near native (TSR_2_), and an unbound state.

In particular, we identify the structural parameter *S* and reorientational decay times *τ* as in our previous study [49] (See methods and SI) for interfacial waters in the three states taken from a reactive dissociation path from the 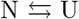 transition path ensemble. We run short NVE MD simulations at 330 K and 300 K and analyse the water structure and dynamics. The results are qualitatively the same at 300 K and 330K (see Figure. 10 and Figure. S19, respectively). In the native state water around the native contact region – comprising residues D33, H146, I147, R148, L149, S150–exhibits two hydration populations. The first is a fast reorienting and more tetrahedrally structured (*S >* 0.4 and *τ <* 4) water population, labelled *tetrahedral fast*, The second is a slowly reorienting, less structured water population denoted *disordered slow* (*S <* 0.4 and *τ >* 4). Note that the bulk water structure parameter is S= 0.38 at 300 K, substantially lower than the tetrahedral fast population. Upon protein association the tetrahedral fast water population increases, as *S* shifts from 0.4 to 0.5 (Figure. 10a,b,c). Since the tetrahedral fast water population lives near hydrophobic groups, we computed the *S − τ* correlation plot for water around the hydrophobic amino acids I147 and L149 (Figure. 10d,e,f). Although these hydrophobic amino acids do not exhibit much tetrahedral water structure in the unbound state, they do so in the native interface where these amino acids are opposite to each other. Moreover, these hydrophobic amino-acids also show substantial disordered slow water populations, due to the influence of the hydration state of the neighbouring charged H146, R148 and (one of the) D33 residues. In contrast, some amino-acids, such as the charged residue D33, exhibit only the disordered hydration state in the N and the U state (Figure. 10g,h,i). Indeed, the slow population around D33 increases upon association as the formation of the hydrophilic interface provides polar and charged neighbouring amino-acids, e.g R40, as well as a more excluded volume (dry interface) environment [48]. However, in TSR2, there is a temporary increase of fast tetrahedral water, because the second D33 has not (yet) formed a contact with R40, but is exposed to hydrophobic residues I147, I29 and L149. Moreover, the smaller dry area of TSR2 compared to N excludes the solvent around the second D33 less, yielding a faster reorientation dynamics and more tetrahedral structure [48].

**FIG. 10:**
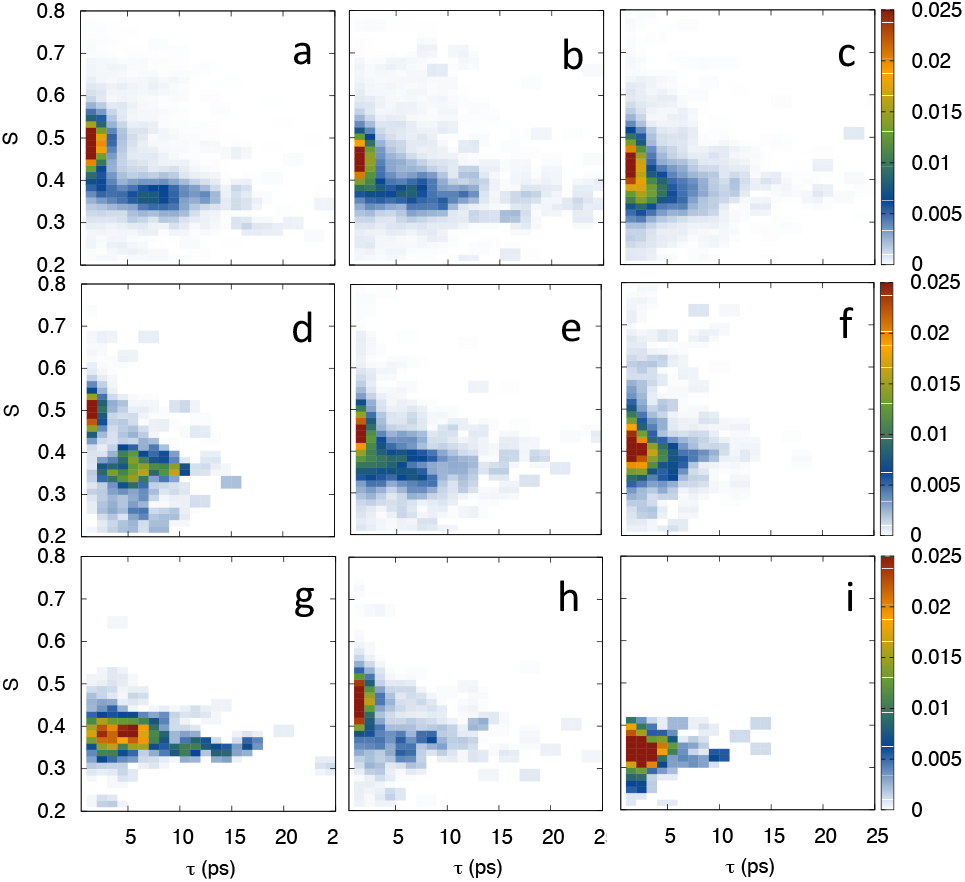
Two dimensional histograms of the reorientation time *τ* versus structural parameter *S* or water molecules residing at the native interface from NVE simulations performed in the a) native, b) transient near native (TSR_2_) and c) unbound states at 300 K. Structural parameter-reorientation time correlation for the native state hydrophobic residues I147, L149 at d) native e) near native and f) unbound configurations. Structure reorientation correlation for the hydrophilic residue D33 at g) native, h) near native, and i) unbound configurations.

Native contacts are supported by water mediated interactions: bridging waters hydrogen bonded to both proteins. In Figure. 11 we plot the hydrogen bond bridge survival correlation function, which decays slower for waters in the hydrophilic contact-rich native state interface compared to waters in the hydrophilic contact-poorer near-native state (TSR2). The faster decay in the transition state indicates proteins are more mobile, as expected. The slow decay in the native state reflects the presence of long lived disordered water around charged and polar amino-acids in the interface. Indeed, the average residence time of the disordered water population increases upon binding, namely 36 ps in the unbound state and 69 ps in the native state, which is much longer than that of the tetrahedral waters, with residence times 17.2 ps and 16.3 respectively (see Figure. 11b)

**FIG. 11:**
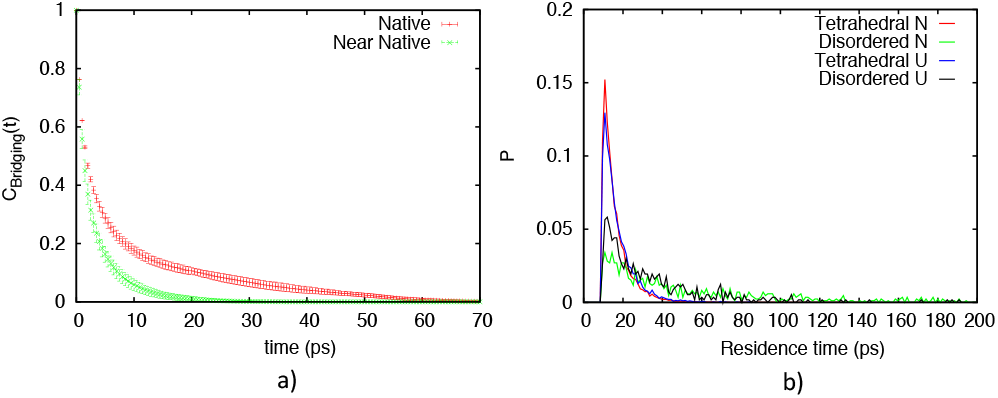
a) Hydrogen bond bridge survival correlation function between any amino acid pair of the interface for native (red) and transient near native states (green) at 300 K. b) Distribution of residence times for the tetrahedral and disordered water population.

Thus, water at the native dimer interface consist of a *disordered slow* population due to formation of long-lived hydrogen bonded bridging water, and a *tetrahedral fast* population, reorienting faster than bulk, which is typical of hydrophobic solvation and thus characteristic of the *β*-lac’s mixed polar-apolar interface. This conclusion also extends to water in the transition state, which has more mobile hydrogen bond bridging waters enabling enhanced protein rotation. While the *β*-lac protein dimer is not in the electrostatically steered regime, the slow reorientation of the disordered slow waters could be related to the reduction in dielectric permittivity as observed by Ahmad *et al.* [12] for the Barnase-Barstar complex. Indeed, this permittivity was found to be small even at larger distances, indicating small fluctuations in the dipole moment, in turn implying slower water reorientation dynamics [12].

As the TPS paths were performed at 330 K we also analysed the water structure and dynamics for this elevated temperature. To check whether lowering the temperature to room temperature would severely alter the results, we repeated the analysis for 300 K. Our findings indicate that the hydration states of water do not change qualitatively upon lowering the temperature from 330 K to 300 K (see Figures. S15-S17).

## CONCLUSIONS

In this study we performed extensive TPS simulations of specific and non-specific dissociation of the hydrophilic *β*-lactoglobulin dimer. This resulted in ensembles of unbiased dynamical transition paths that are inaccessible with standard MD. Analysis of these sampled path ensembles revealed that specific dissociation can occur either via a direct aligned transition, a hopping and re-binding transition followed by unbinding, or via a sliding transition before unbinding.

For non-specific dissociation, the trajectories are significantly shorter, but also exhibit mechanisms involving *direct* dissociation, a *sliding* mechanism where first the dry area decreases by a sliding movement, before dissociation, and a *hopping* dissociation route where the proteins first rebind before completely dissociating. This finding suggests that the mechanism of direct dissociation, sliding and rebinding are general dissociation mechanisms. Indeed, theoretical and numerical work on simple models shows that indeed protein association is influenced by rebinding [25, 26]. The sliding mechanism can thus be viewed as rebinding to a non-specific state without full solvation.

Employing reaction coordinate and transition state analysis we found that in the transition states regions only a small fraction (~ 25%) of the native contacts are present. This conclusion is in agreement with recent straightforward simulation by the DE Shaw group [19]. In addition, we found evidence for an important role of the D33-R40 salt-bridge, also implicated by experiments[29]. Moreover, we investigated the role of the solvent in the dissociation process by assessing the structure and dynamics of the solvent molecules. This analysis revealed that the dry native interface induces enhanced populations of both disordered hydration water and hydration water with higher tetrahedrality, mainly nearby hydrophobic residues. We conclude that water assists (un)binding through formation of a transient complex [24] characterised by hydrogen bond bridging waters, thus facilitating rebinding[23].

In summary, the rare unbiased reactive molecular dynamics trajectories shows in full detail how proteins can dissociate via complex pathways including (multiple) re-binding events. Our results give an unbiased dynamical view of the mechanism of protein-protein dissociation in explicit solvent, as well as insight in the structural and dynamical role of the solvent in this process. The atomistic insight obtained assists in further understanding and control of the dynamics of protein-protein interaction including the role of solvent. We expect that our predictions can be experimentally tested, e.g. with spectroscopic techniques such as NMR, or vibration sum frequency generation.

Finally, we remark that our approach does not provide a complete full sampling of the dissociation or, because of the time reversible nature of the dynamics, the association process. The main reason for this is the limited allowed duration of the pathways. Restricting the paths to a maximum time excludes all paths that show many hoppings between intermediates. On the other hand the fact that there is still a substantial number of successful dissociation paths accepted shows that these long re-binding paths might be not so important for dissociation. Nevertheless, we stress that for a full kinetic description of the entire dissociation and association process much more sampling is needed. A viable way might be to combine the path sampling approach with the MSM methods of Ref. [17].

## ACKNOWLEDGEMENT

The research leading to these conclusions was funded by the NanoNextNL Programme, a micro and nanotechnology consortium of the Government of the Netherlands and 130 partners. We acknowledge support from the Nederlandse Organisatie voor Wetenschappelijk Onderzoek (NWO) for the use of supercomputer facilities.

